# Sequence mismatch between gene-drive and target-site flanking regions significantly impairs homing efficiency in *Culex quinquefasciatus*

**DOI:** 10.1101/2025.08.01.668174

**Authors:** T. Harvey-Samuel, R. Kaur, P.T. Leftwich, X. Feng, V. Gantz, L. Alphey

## Abstract

CRISPR/Cas9-based homing gene-drives (homing-drives) hold enormous potential as control tools for mosquito disease-vectors. These genomically-encoded technologies spread themselves through target populations by creating double-stranded DNA breaks on homologous chromosomes, into which the homing-drives are copied (‘homed’). Homing is dependent on sequence homology between the genomic regions flanking the transgene insertion and the break site. Homing efficiency (i.e. copying rate) substantially impacts the power of these systems: less efficient homing-drives spread slower, have fewer applications and are more resistance-prone. Understanding what influences homing-drive efficiency is therefore vital to the successful use of these technologies. Here we report a novel mechanism by which a homing-drive’s efficiency can be significantly impaired by natural sequence variation within a population into which it is spreading. Using a *kmo*-targeting ‘split’ homing-drive in the West Nile virus mosquito *Culex quinquefasciatus*, we found that target-site heterology (sequence mismatch between the genomic regions flanking the target cut-site and the homing-drive transgene) of less than 10% reduced homing efficiency by up to 54%. While substantial research effort has been dedicated to increasing homing-drive efficiency through optimisation of within-construct components, our results highlight that the real-world efficacy of these systems may in part depend on variation beyond these controllable factors.

## Introduction

CRISPR/Cas9 ‘homing’ gene-drives (henceforth homing-drives) are genomically encoded technologies which hold immense promise for controlling intractable pests, with notable development in disease-vectoring mosquitoes(1–7). A homing-drive increases its allele frequency from one generation to the next by copying (‘homing’) itself from one homologous chromosome to another in germline cells (8). For pest management, a homing-drive can be designed to simultaneously spread traits beneficial for control e.g. pathogen refractoriness or a genetic load for population suppression/eradication (9). The efficiency of the homing reaction substantially influences the population-level efficacy of a homing-drive: higher homing rates enable faster spread and higher tolerance of drive-associated fitness costs (10).

At a molecular level, homing occurs when Cas9 expressed by a homing-drive transgene creates a double-stranded DNA break (henceforth ‘cut-site’) at a specific sequence on the target chromosome, into which the homing-drive is homed (11) (Fig 1.). Homing is mediated by the homology-directed-repair (HDR) pathway, requiring the genomic sequences flanking the homing-drive transgene insertion-site and the target allele cut-site to show homology. While substantial effort has been devoted to exploring how ‘within-drive’ components (e.g. regulatory elements for expressing Cas9 (12,13)) affect homing rates, comparatively little attention has been paid to the role of these genomic flanking regions in influencing homing efficiency. This is surprising as the impact of even minor sequence divergence between loci in substantially reducing homologous recombination rates is well established (14). In the real-world, homing-drives will be required to function in genetically diverse ‘wild’ populations, which will likely also include pre-existing heterogeneity around the cut-site. Whether sequence mismatch (henceforth ‘heterology’) between a homing-drive’s flanking genomic regions and those flanking its target allele cut-site influences homing rates is thus of considerable importance.

**Figure 1:**
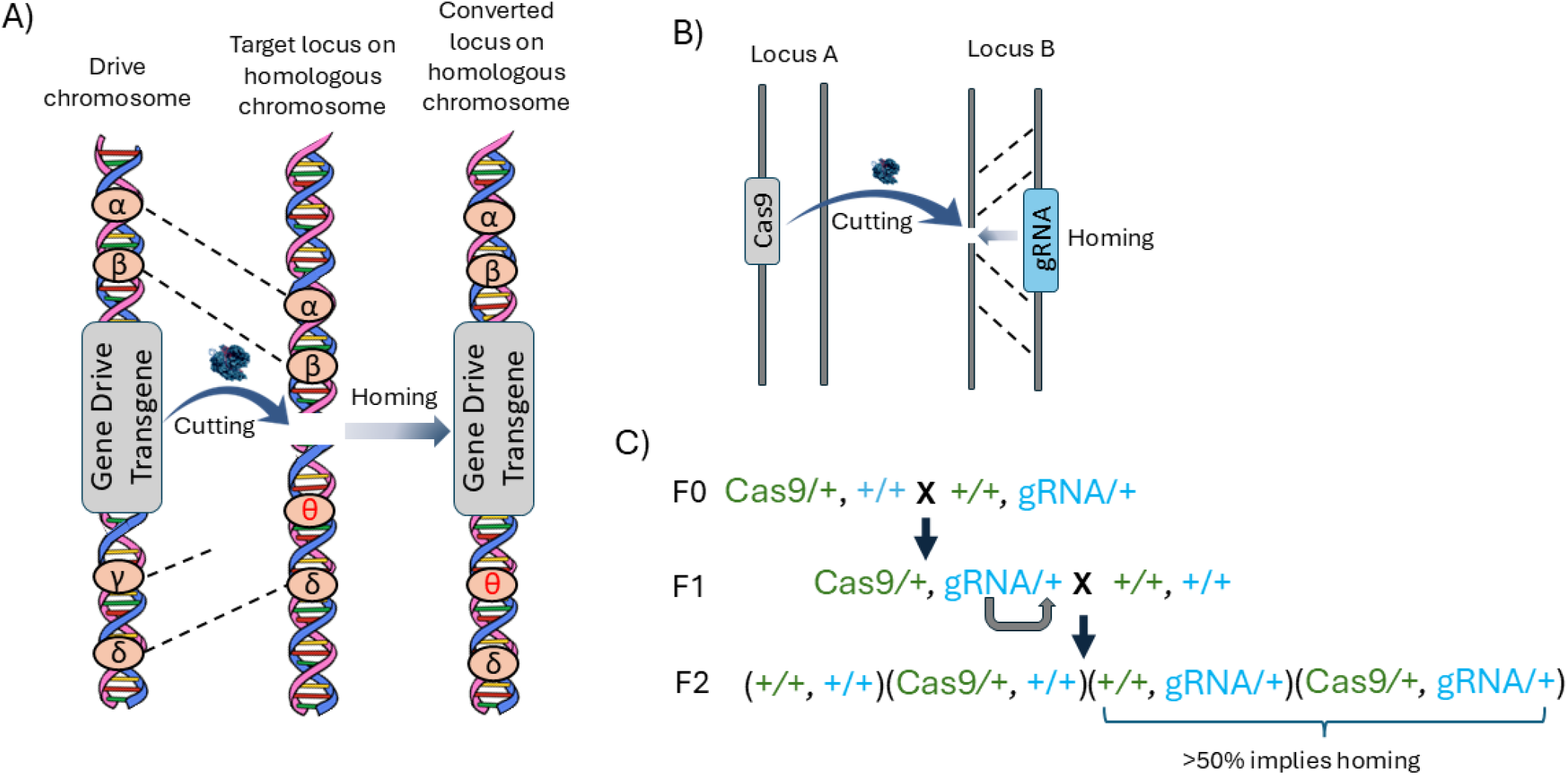
Schematic representations of CRISPR-Cas9 based homing gene-drive technologies. A) A ‘global’ homing gene-drive in which all the molecular components required to initiate the homing reaction are linked and integrated at a single locus in the genome which conforms to the target site of the integrated gRNA. Expression of the Cas9 enzyme and gRNA thus mediates the creation of a double-stranded break (DSB) at the insertion locus on the homologous chromosome. This DSB can be repaired through the Homology-Dependent Repair (HDR) pathway which results in the copying of the gene-drive transgene onto the homologous chromosome. If this process occurs in germline cells, it will result in super-mendelian inheritance of the gene-drive. The efficiency of HDR is dependent on homology between the sequence used as the repair template and the sequences flanking the DSB. These sequences, however, are not always perfectly matched, as shown in the schematic (specifically, γ and θ flanking sequences show heterology). B) Schematic representation of a ‘split’ homing gene-drive, here where the gRNA-cassette is located within the homing locus (Locus B) as with a global gene-drive design, but the Cas9-cassette is now located at an unlinked, non-homing, location (Locus A). C) The mosquito line crossing scheme for combining the Cas9-transgene and the gRNA-transgene in a split-drive system. In the ‘grandparental’ F0 generation, the two lines are crossed. In the ‘parental’ F1 generation, individuals carrying both transgenes (trans-heterozygotes) are isolated and crossed to wild type, producing the F2 generation where transgene fluorescence inheritance ratios are recorded in order to estimate homing rates. In this representation, Locus B (i.e. that containing the gRNA cassette, aka the homing element) is coloured in blue, while Locus A (i.e. that containing the Cas9 cassette) is coloured in green.

We explored this question using our homing-drive system in the West-Nile-virus vector mosquito *Culex quinquefasciatus* (5). This system consists of a ‘split-drive’ design where the two homing-drive components (germline-expressing Cas9 and gRNA-expressing cassettes) are integrated at independent loci - one of which (here the gRNA-cassette) is in the target allele - and maintained in separate lines (11). The two lines can then be crossed together to initiate the homing reaction in trans-heterozygotes. By using the *kynurinine 3-monooxygenase* (*kmo*) gene as our target allele (where homozygous null mutations give an easily visualized ‘white-eye’ phenotype (15)) and conducting all comparison crosses in a ‘controlled’ genetic background, we were able to unambiguously distinguish Cas9-mediated cutting and homing, and attribute variation in these to individual *kmo* target alleles which displayed varying levels of heterology. This experimental design provided unparalleled power to assess this fundamentally important question.

## Results and Discussion

Our aim was to investigate the effect of varying the heterology of sequences flanking the cut-site of a target allele on a homing-drive’s homing efficiency. We assessed this at two distinct levels, each specifically designed to give insight into the behavior of this potential effect. Firstly, we took an ‘allele-by-allele’ approach – assessing a homing-drive’s efficiency as it was paired against four different *kmo* target alleles in a controlled genetic background. Secondly, we extended this analysis to a broader range of cross schemes and sexes to assess the generality of findings.

### 1: Allele-by-allele approach

In ‘split’ homing-drives, one of the expression cassettes (here, that expressing the gRNA) is integrated into the target allele at the precise cut-site, forming the ‘homing-element’. This can then be combined with the other element, here encoding Cas9, to provide trans-heterozygotes (aka “double heterozygotes”) which have one copy each of Cas9 and homing-element and so can catalyse the homing reaction. Furthermore, and crucially for our purpose, in this set-up, the target allele into which a homing-element attempts to home will always come from the line (and therefore the genetic background) which contributed the Cas9 transgene. Preliminary analysis of our highly inbred wild-type (WT) line (in which the *kmo*-gRNA line was generated and is outcrossed to each generation – TPRI genetic background – see methods) and the *Vasa*-Cas9 line (which was generated separately in the CA genetic background – see methods) showed that these two lines possessed highly divergent *kmo* allele sequences. Those carrying the TPRI genetic background contained only a single *kmo* sequence (allele A) which perfectly matched the sequences flanking the integrated *kmo*-gRNA homing-element, while the *Vasa*-Cas9 line contained three different *kmo* allele sequences (alleles B, C and D) (See figure 2A). We took advantage of this to design an assay in which a Cas9-bearing individual could contribute a variety of target alleles to a homing cross, some but not all of which would match the homing-element flanking sequences but, critically, all of which would end up in the same mixed genetic background (see figure 2B). These female trans-heterozygotes (*kmo*-gRNA/+ ^A,B,C or D^, *Vasa*-Cas9/+) were then crossed to male *kmo* -/- homozygotes to allow us to detect both homing (deviation in inheritance rate of homing-element from Mendelian expectation of 50%) and the rate of end-joining (individuals inheriting loss-of-function *kmo* alleles from their mother, generated via Cas9 cutting followed by error-prone repair, will have white eyes but lack the fluorescent marker of the homing-element) in their progeny. This assay showed a highly significant effect of the *kmo* target allele (i.e. A,B,C or D) on homing rate (LRT: χ^2^_3_ = 16.17, p = 0.001), with a 54% difference in estimated homing rates associated with the highest (A = 0.37 [95% CI; 0.31 – 0.42]) and the lowest (D = 0.17 [0.04 – 0.29]) *kmo* target alleles (Figure 3, Supplementary Figure 1, Supplementary Table 1, Supplementary Table 2). Preliminary sequencing confirmed that the gRNA target sequence was present in all four *kmo* alleles and this was confirmed by our comparison of estimated cutting rates (i.e. estimated homing + observed end-joining) between the *kmo* alleles, which did not significantly differ (LRT: χ^2^_3_ = 5.27, p = 0.15; Supplementary Figure 1, Supplementary Table 1, Supplementary Table 2). This suggests that the differences in homing we observed between the four target *kmo* alleles were not simply driven by Cas9 being able to cut some alleles more efficiently than others. Instead, taken together, these data strongly suggest that the mechanism underlying the observed reduction in homing is impairment of homology-directed repair through drive-target allele flanking sequence mismatch.

**Figure 2:**
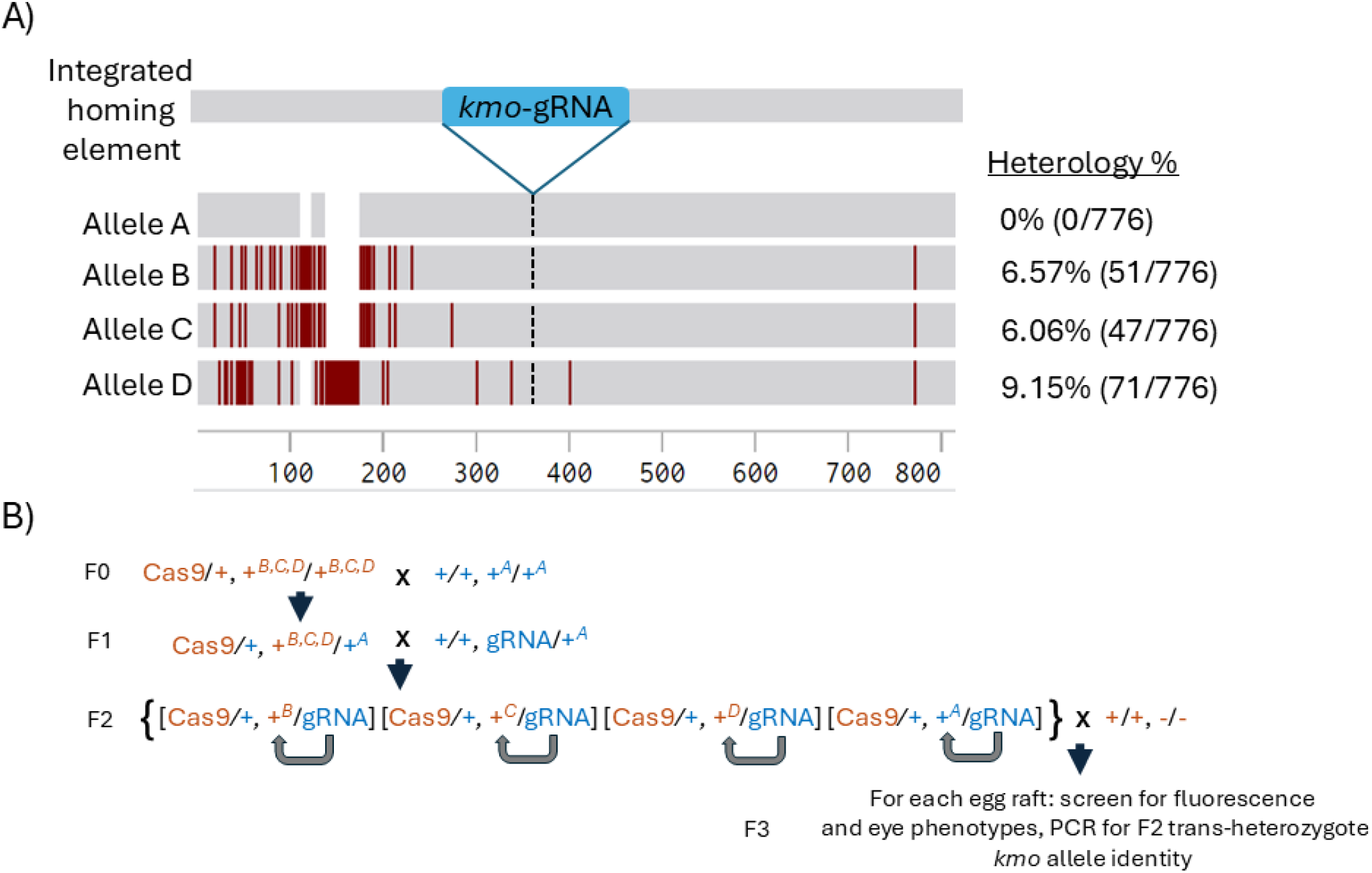
Details of the *kmo* locus sequence targeted and genetic background of lines crossed in Assay 1 (allele-by-allele approach). A) Representations of the four *kmo* alleles identified through Sanger sequencing. Allele A was the only allele observed in the ‘TPRI’ genetic background and showed 100% homology over the sequenced region to the *kmo*-gRNA integrated homing element (0 mismatches over 776bp). Alleles B, C and D were the only alleles observed in the ‘CA’ genetic background and showed varying levels of heterology over the sequenced region to the *kmo*-gRNA integrated homing element. B) Schematic showing how introgression of the two genetic backgrounds allowed investigation of the influence of target allele identity on homing rates. In brief, in the F0 generation, individuals carrying the *Vasa*-Cas9 transgene (which resides in the CA genetic background – represented in orange) were first crossed to wild-type individuals possessing a TPRI genetic background (represented in blue). *Vasa*-Cas9 heterozygotes, now in a hybrid CA/TPRI genetic background were then crossed to individuals from the *kmo*-gRNA line (which possesses a TPRI genetic background) (F1 generation). This produced the F2 generation where the *kmo-gRNA* cassette was present opposite all four identified *kmo* alleles, in proportion to their frequencies in the F1 *Vasa*-Cas9 heterozygotes. This allowed the homing reaction to take place into four separate target alleles, at the same locus, in a controlled genetic background (shown by curved arrows). In this generation, trans-heterozygotes were crossed to individuals from a *kmo* -/- strain (CA genetic background). This produced the F3 generation where, for each egg raft, fluorescence ratios and eye phenotypes were recorded and the target *kmo* allele present In the F2 trans-heterozygote which produced that egg raft identified through PCR.

**Figure 3:**
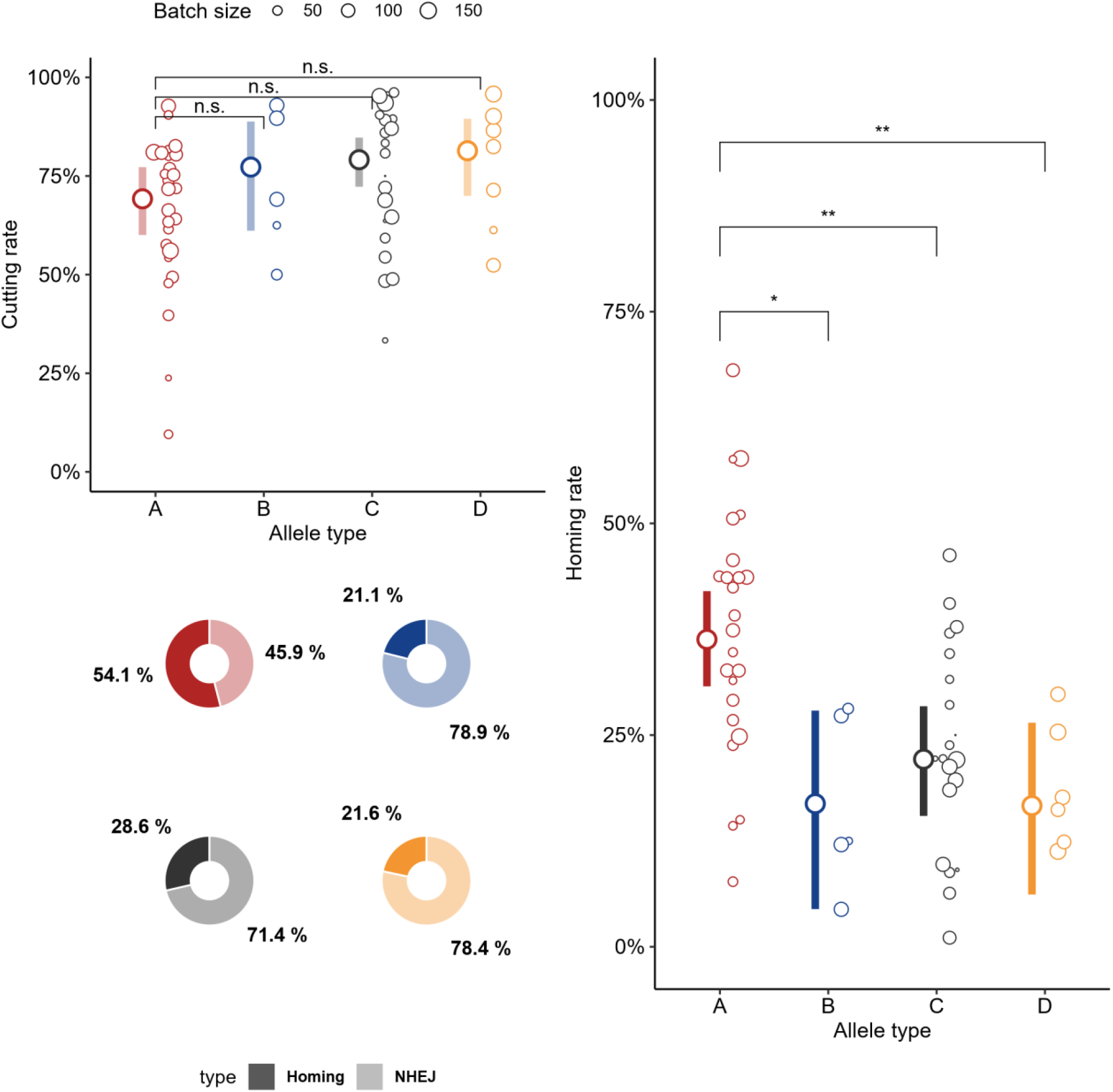
Results from Assay 1 homing crosses: A) ‘Cutting rates’ estimated for the four identified *kmo* target alleles in the F2 *Vasa*-Cas9/*kmo*-gRNA trans-heterozygotes. Here cutting was calculated as the sum of NHEJ (signified by white-eye phenotype individuals) and HDR (derived from *kmo*-gRNA inheritance >50%) events observed in the F3 progeny. B) ‘Homing rates’ estimated for the four identified *kmo* target alleles in the F2 *Vasa*-Cas9/*kmo*-gRNA trans-heterozygotes. Here homing rate was calculated as the % of wild-type alleles converted to bear the homing-drive cassette in the trans-heterozygote germline (derived from level of *kmo*-gRNA inheritance >50%). As such, a homing rate of 50% would represent half the wild-type alleles in the F2 trans-heterozygote germline being converted to homing-element alleles, and an overall inheritance rate of the *kmo*-gRNA transgene of 75%. For both A) and B) Large symbols and error bars (vertical lines) represent mean and 95% confidence intervals calculated by a generalized linear mixed model with a binomial (‘logit’ link) error distribution, individual points to the right of each estimated mean represent pools of offspring derived from a single female parent. Relative size of the small points is in proportion to the number of individuals recorded for that data point (batch size). C) Raw data values of recorded ‘Homing rates’ as a percentage of the pooled total of ‘Cutting rates’ for each of the four identified *kmo* target alleles. Raw data proportions (C) represent unadjusted observations, while model-estimated means (A &B) account for covariates and random effects.

### 2: Generalised approach

In the real world, homing-drive transgenes could be inherited from, and be active in, either sex. Previous work has shown that the efficiencies and dynamics of homing-drives can vary dramatically dependent on direction of inheritance (paternal/maternal) in parents and grandparents (e.g. (4,16,17)). As such, we were interested in investigating the extent to which the observed influence of heterology on homing rates was maintained when a wider variety of cross combinations were considered. To achieve this, we first introgressed the *Vasa*-Cas9 transgene into the TPRI genetic background by outcrossing *Vasa*-Cas9 individuals to our TPRI wild-type line for three consecutive generations. This TPRI introgressed *Vasa*-Cas9 line (henceforth *Vasa*-Cas9^i^) was then crossed to the *kmo*-gRNA line in a full factorial homing-drive cross (i.e. where the *Vasa*-Cas9 and *kmo*-gRNA transgenes were inherited from both grandparental (F0) sexes and the trans-heterozygous (F1) individuals were of either sex) and results compared to our previously published homing analysis of these two transgenes - in that case where the *Vasa*-Cas9 transgene was in a pure CA genetic background (5). We found that in all like-for-like comparisons the homing rates in this study were higher than in our previous study (aggregate difference 57.8% - 69.3%), only in the male-male crosses was this increase not statistically significant (Figure 4, Supplementary Table 3). These results further support our conclusion that target-site/homing-element flanking-sequence heterology can significantly affect homing efficiency.

**Figure 4:**
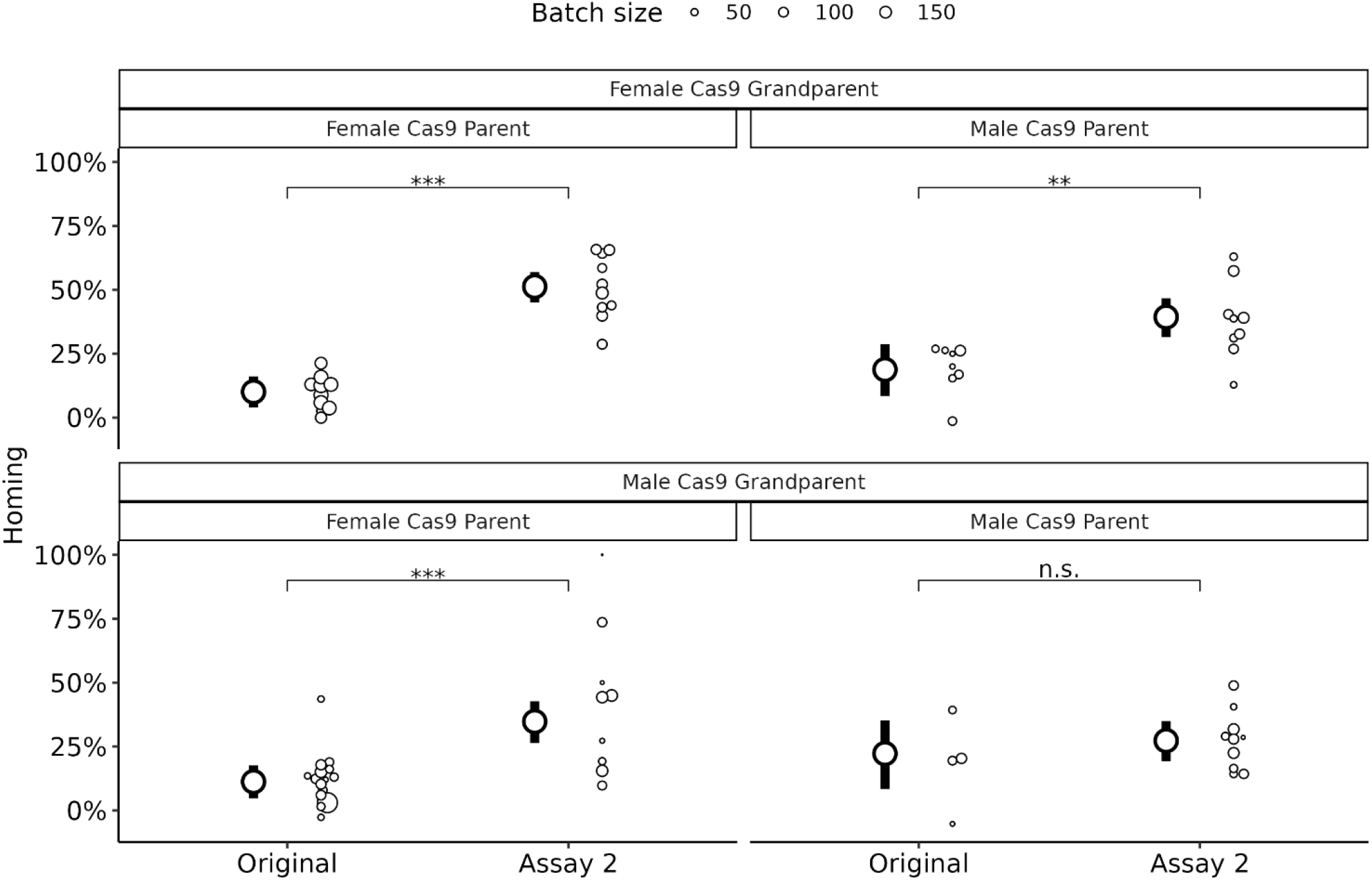
Comparison of estimated homing rates from (5) ‘Original’ (i.e. Harvey-Samuel & Feng, 2023) and Assay 2 (i.e. Generalised approach - this study). In each case, a full factorial split-drive cross setup was applied (i.e. testing all combinations of grandparental and parental sex on estimated homing rates). Facets separate offspring according to the grandparent and parent from which they inherited the *Vasa*-Cas9 transgene. Large symbols and error bars represent mean and 95% confidence intervals, small points represent raw data. Relative size of the small points is in proportion to the number of individuals recorded for this data point (batch size). Annotated asterisks represent pairwise significant difference tests at (n.s. = non-significant, * <0.05, **<0.01, ***<0.001).

To date, only one other study has directly explored the influence of significant (i.e. >1%) flanking-sequence heterology on homing-drive performance (in *Anopheles gambiae*) (18). In contrast to the results presented here, that study observed no significant influence of target-site heterology on homing-efficacy, despite similar levels of sequence divergence (5.3-6.6% over c.690bp). What might explain the differences between this finding and ours? We propose three non-exclusive possibilities.

The first involves the genetic backgrounds in which these crosses were conducted. Pescod *et al*. (18)crossed their homing-drive directly into different geographically distinct *An. gambiae* strains, each of which contained divergent target alleles. As such, each of the different homing-drive/target allele assessments occurred in a distinct genetic background. As it has been shown that (as with ‘natural’ homologous recombination(19)) many loci unlinked to a homing-drive may exert significant influence over homing efficiencies (20), this experimental design may have introduced additional variation, potentially obscuring the effects as observed here. We were able to control for these potential epistatic effects by crossing, and therefore partially introgressing, our *Vasa*-Cas9 line into the TPRI background prior to homing assays, ensuring each of our *kmo* target allele comparisons occurred within (on average) the same, mixed CA/TPRI genetic background.

Secondly, there are potentially species-specific explanations for these contrasting results. Homing-drive efficiencies in *Anopheline* mosquitoes are uniquely high (often approaching or reaching 100%) (e.g. (21)). The precise reason for this is unknown but could include less stringent requirements on engaging the HDR pathway e.g. higher tolerance to flanking-sequence heterology, or smaller flanking sequences required to initiate homing. Indeed, analysis of homing conversion tract lengths in *An. gambiae* (i.e the degree to which the flanking regions of a target allele are converted to carry heterologous sequences contained within the homing-drive flanking regions) showed that (beyond the homing-drive cassette itself) only small sequences are transferred onto the target chromosome (>80% of conversion tracts <50bp)(22). This is substantially shorter than reported for studies considering ‘natural’ homologous recombination in other species (typically c. 200bp (19)) and may signify shorter required sequences for RAD51-dependent complementary strand recognition/invasion in *Anophelines*. In contrast, conversion tract lengths in *Aedes aegypti*, another Culicine mosquito, following CRISPR-Cas9 plasmid-based knock-in (also dependent on HDR, though likely occurring in somatic rather than germline cells) were substantially longer (mean of 160bp) (23). Interestingly, and in concurrence with our results, that study in *Ae. aegypti* also identified significant negative effects of plasmid/target-site heterology on the efficiency of knock-in. Taken together, these results could suggest that the HDR-pathway in *An. gambiae* is simply less affected by sequence heterology than in other species, including Culicine mosquitoes.

Finally, the near-100% homing rates displayed by the drives tested in Pescod *et al* may have made it difficult to observe anything other than extremely large changes (i.e. reductions) in homing, potentially underestimating or missing ‘subtler’ modifiers of homing-efficiency – e.g. those caused by target alleles less amenable to homing. This is because in a homing-drive, inheritance-bias (i.e. level of homing) is described by odds (p/(1-p)) – where p = wild-type allele conversion probability - and the effect of a change in odds on the change in homing rate differs substantially depending on the initial level of homing ((10), Figure 5). In essence, for homing-drives which display very high homing-rates, even large changes in odds will have only small effects on the observed levels of homing. This effect is much less pronounced in less efficient homing-drives, like that demonstrated here. The diminishing return on changes in odds means that, near 100% efficiency, subtle effects on homing may go unnoticed, complicating efforts to accurately evaluate or identify modifiers of highly effective homing-drives.

**Figure 5:**
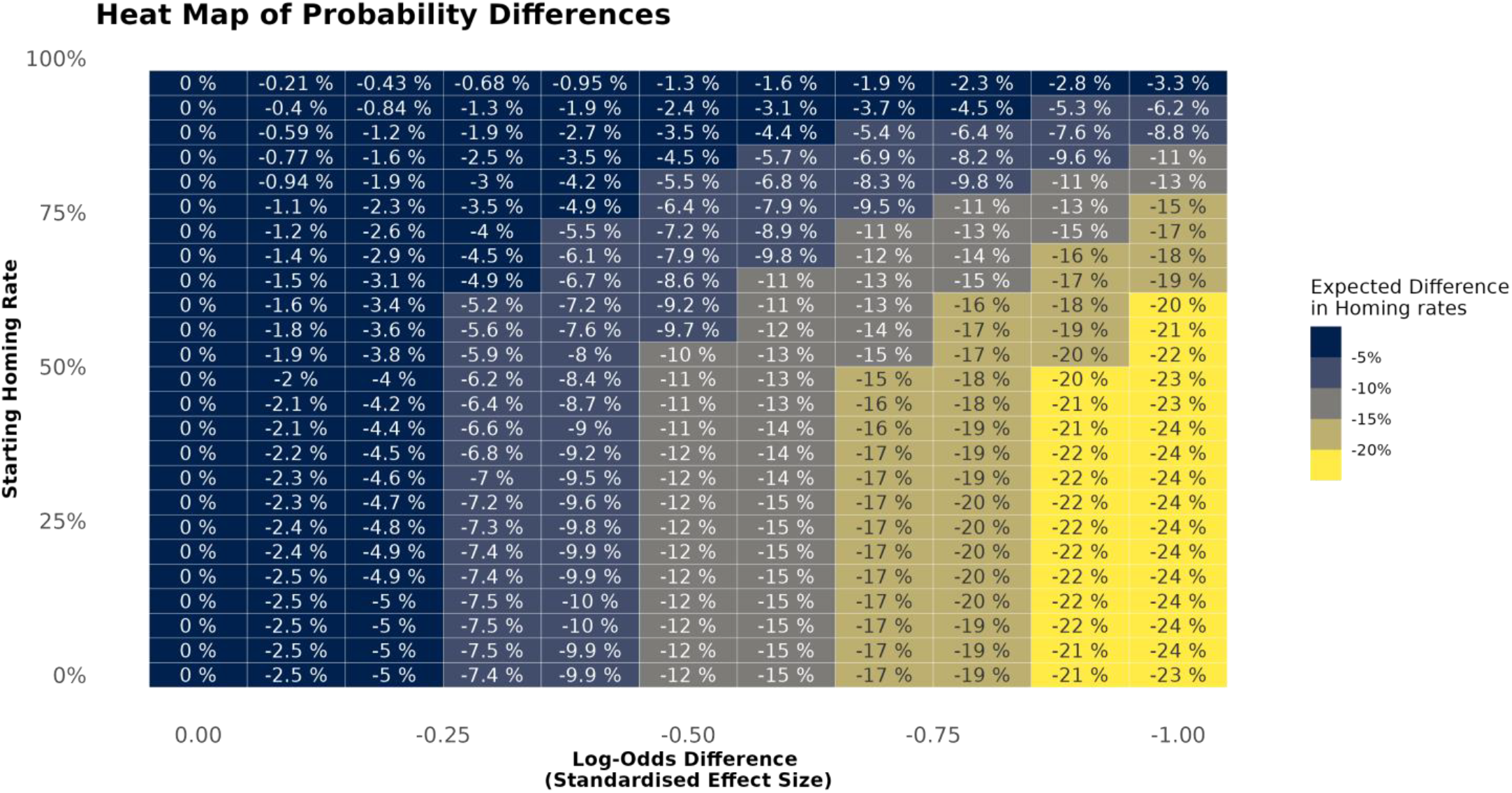
Variation in observed probability change (e.g. change in homing) based on starting probability and Log-Odds effect size (e.g. change in odds of alle conversion) This heatmap illustrates how a standardized change in log-odds affects observed probability differences, depending on the initial probability level. The y-axis represents the starting probability of an event (here the level of homing), while the x-axis indicates the standardized effect size of a change in log-odds. The color gradient reflects the magnitude of the resulting probability difference, with lighter regions signifying larger changes. The effect of the same log-odds shift is most pronounced when initial probabilities of an event are in the mid-range (e.g. around 50%, i.e in the context of inheritance-bias - where homing is calculated at 0%) and diminishes as probabilities approach the extremes (near 100% inheritance/homing). This highlights the non-linear nature of probability changes in a logit model, where the same predictor shift has varying impacts depending on the starting likelihood of an event.

In conclusion, our results demonstrate a novel factor affecting homing-drive efficiency. We show that heterology between a homing-drive’s flanking regions and those sequences flanking the target allele cut-site can significantly reduce that drive’s performance. A critical question is to what extent this effect could impact homing-drive performance in the real-world. In most cases it should be assumed that homing-drives could encounter significant levels of sequence variation upon release, particularly in continental areas where most of their deployments will likely occur. In our system, even the most divergent *kmo* allele tested was still able to be targeted for Cas9 cleavage and successfully converted to carry the homing-drive, albeit at a significantly reduced rate. If the percentage reduction in homing efficiency observed here were to translate to released strains, however, this could indeed substantially impede their spread through a target population (Supplementary Figure 2). While naturally high homing rates may mitigate this effect, for the vast majority of species which have been tested so far, reported homing rates are modest (5,7,24–26) and thus they may be, to a degree, affected by this mechanism. Prior knowledge of these effects for a given species, homing-drive locus and target population may allow future release programs to preempt adverse consequences on project outcomes, e.g. by increasing release rates accordingly to counteract predicted lower drive efficiency, or by introgressing the homing-drive into a particularly homing-recalcitrant allele ahead of releases. Additionally, our findings highlight the utility of exploring factors which affect homing-drive dynamics and performance in species/model-systems which do not display near-perfect performance. While ‘high-performance’ homing-drives may be the first in line for field-release, it is precisely their extreme efficiency which may make them less useful for investigating some of the underlying factors which affect the drive mechanism itself.

Our results are particularly significant as gene-drives edge ever closer to field application. They suggest that, at least in some cases, homing-drive efficiencies estimated in genetically homogenous lab populations may be unrepresentative of those which would occur in the real-world. While lab-field discrepancies are well known for other parameters relevant to genetic biocontrol, e.g. mating success, longevity or other factors relating to fitness, this is the first study we are aware of which identifies such a disparity related to the homing mechanism itself. This information will be critical in helping to set realistic expectations of these potentially game-changing systems, if and when they enter the field-testing phase.

## Materials and methods

### Ethics

Work followed procedures/protocols approved by The Pirbright Institute Biological Agents & Genetic Modification Safety Committee. All homing assays were conducted at The Pirbright Institute IS4L arthropod containment facility under the necessary safety regulations for gene-drive research.

### Mosquito lines used, rearing and maintenance

The *kmo-*gRNA line was generated previously by integrating a *kmo*-gRNA expression cassette via CRISPR/Cas9 HDR into the *kmo* gene in the TPRI (Tropical Pesticides Research Institute) genetic background. The *kmo* -/- line was generated previously by CRISPR/Cas9 knockout of the *kmo* gene in the CA (California) genetic background. The *Vasa*-Cas9 line was generated previously by integrating a Cas9 ORF under the transcriptional control of *Cx. quinquefasciatus Vasa* regulatory elements into the *Cardinal* gene in the CA genetic background via CRISPR/Cas9 HDR. The Wild-type (WT) line is the unmodified TPRI line. Rearing/maintenance followed procedures previously reported.

### Mosquito experiments

#### Assay 1: Allele-by-allele approach

Homozygous *Vasa*-Cas9 females were first pool-crossed to WT males, producing heterozygous *Vasa*-Cas9 F1 progeny in a 50:50 TPRI/CA genetic background. Female F1 were then pool-crossed to heterozygous *kmo*-gRNA males to give the F2 generation. Trans-heterozygous *Vasa*-Cas9, *kmo*-gRNA female F2 progeny were then pool-crossed to *kmo*-/- males and F3 egg-rafts produced individually isolated, with each egg-raft being from an individual female trans-heterozygote. L3-4 stage progeny from each egg-raft were scored for presence of transgenes/eye-pigmentation (white-eyes), as previously demonstrated.

#### Genotyping of target allele (Assay 1 only)

After scoring, gDNA was extracted from a single WT or *Vasa*-Cas9 individual from each F3 egg-raft (Machery-Nagel Nucleospin Tissue kit, Düren, Germany). This was used to PCR-genotype the maternally-contributed *kmo* allele (i.e. the allele into which the *kmo*-gRNA had attempted to home in the previous generation). The paternally-contributed *kmo* allele was excluded by designing the reverse primer (PL506) to lie across a region deleted during the generation of the *kmo* -/- line. The primers were designed to bind to all 4 *kmo* alleles identified in a preliminary analysis of the strains utilized (A,B,C,D). PCR conditions as follows 98C-1min, (98C-30s, 72C-15s, 72C-1min) x 35, 72C-2min. 200ng of gDNA used per 50ul Q5 PCR Reaction (New England Biolabs). Primers = F(PL303): CCAACATTCACCTTCACTTCAACCACAAGC and R(PL506): GTGAGCGTCCTCCCACCGAG. Amplicons extended c.320/540bp upstream/downstream of the cut site – sufficient distance to distinguish the 4 alleles. Reactions were run on a 1% agarose gel and bands extracted using the Machery-nagel Nucleospin Gel and PCR clean-up kit. Purified bands were Sanger sequenced using the forward primer PL303 and sequenced amplicons aligned to the TPRI *kmo* sequence.

#### Calculation of homing and cutting rates (Assay 1 only)

Homing rates are defined as the percentage of homologous wild-type chromosomes converted to carry the *kmo*-gRNA transgene (i.e. the percentage of WT alleles which have been affected by homing). It was estimated for each egg-raft as (200 x (no. *kmo*-gRNA/total progeny – 0.5). Cutting rate is defined as the percentage of homologous wild-type chromosomes observed as having been cut by Cas9. This includes DSBs that were repaired via HDR (homing), NHEJ (or other error-prone mechanisms) but excludes repairs which resulted in no observable change to the WT allele (perfect repair). It was estimated for each egg-raft as the previously estimated homing rate + (200 x (no. non-*kmo*-gRNA progeny which showed white-eyes/total progeny – 0.5). That is, the HDR + error-prone pathway rates.

#### Assay 2: Generalising findings

Homozygous *Vasa*-Cas9 females were first pool-crossed to WT males, producing heterozygous *Vasa*-Cas9 F1 progeny in a 50:50 TPRI/CA genetic background. This was repeated twice to give F3 *Vasa*-Cas9 progeny in a predominantly TPRI genetic background (c. 87.5% TPRI alleles). Male and female *Vasa*-Cas9 heterozygote F3s were pool-crossed to female and male *kmo*-gRNA line heterozygotes, respectively, giving two cohorts in the F4 generation (one from each of the two F3 crosses). Trans-heterozygous male and female F4 progeny from each of these two crosses were pool-crossed to female and male *kmo* -/- individuals, respectively (4 crosses total), and the egg rafts produced individually isolated and scored as for Assay 1.

#### Statistical analysis

Homing and cutting rates were both analysed with generalized linear mixed models with a binomial (logit) error distribution. Fixed effects for Assay 1 were the genotyped maternally contributed *kmo* allele with individual female egg-batch as a random effect. Minor overdispersion in the cutting analysis models meant all 95% confidence intervals were checked by semi-parametric bootstrapping (1000 iterations). Close agreement between the two sets of intervals indicated minimal overdispersion, suggesting that the model assumptions were adequately met and both summaries are presented. For Assay 2 fixed effects included the sex of the Cas9-bearing parent and grandparent as well as whether the data came from the original (i.e. previously published) experiment or post-introgression (i.e. Assay 2). Individual female egg-batches were also included as a random effect. All models and simulations were constructed in R version 4.3.3 with the package lmerTest, model fits were checked with the package DHARMa and model predictions generated with the package emmeans. Data processing and visualization was performed with the tidyverse packages.

## Author contributions

THS, XF, VG and LA conceived the project. RK, THS and PL contributed to the design of the experiments. THS, RK and PL performed the experiments and contributed to the collection and analysis of data. THS and PL wrote the manuscript. All authors edited the manuscript.

## Data Availability

The authors affirm that all data necessary for confirming the conclusions of the article are present within the article, figures, and tables. Strains and plasmids are available upon request.

